# Measuring magnetic field effects in fluorescent flavoproteins via spin-dependent fluorescence intensity requires photoexcitation to be faster than spin-independent ground state recovery

**DOI:** 10.64898/2026.07.08.737352

**Authors:** Brian L. Ross, Alessandro Lodesani, Clarice D. Aiello

## Abstract

Weak magnetic fields affect many biological processes across the tree of life, though the precise molecular sensors and pathways involved in such magnetoresponses remain mostly uncharacterized. Fluorescence is a useful tool for investigating magnetic field effects in flavoproteins, as their chromophore’s fluorescence intensity can be shown to depend on the spin states of electronic radical pairs. Here, we describe a four-state ordinary differential equation model to understand what parameter sets result in fluorescence contrast between spin states in photocycles with singlet and triplet radical pairs. We conclude that only certain sets of parameters result in the fluorescence intensity being a good proxy measurement for singlet yield. In particular, we observe that the illumination intensity required to obtain fluorescence contrast depends on the rate of the slow spin-independent radical termination reactions that recover ground-state oxidized fluorophores. Moreover, to observe a magnetic field effect in fluorescence intensity when an external magnetic field modulates the singlet yield, the illumination intensity must be strong enough such that photoexcitation is not the rate-limiting step. This understanding suggests that flavoproteins that do not exhibit magnetic field effects in their fluorescence emission under certain experimental setups may still be sensitive to weak magnetic fields in terms of function, as magnetosensitivity in fluorescence depends strongly on illumination conditions.

## 1. Introduction

Decades of research suggest that weak magnetic fields in the *µ*T to mT range can have significant biological effects across a wide array of biological systems and processes [1–5]. One leading hypothesis is that magnetosensitive flavoproteins are key mediators of these effects through a process known as the radical pair mechanism [6]. In this process, electron transfer between an electron donor (aromatic residue or reactive oxygen species) and an excited flavin cofactor (FAD or FMN) generates a spin-correlated radical pair. External magnetic fields can alter the extent of interconversion between singlet (total spin = 0) and triplet (total spin = 1) states for this pair, thereby changing the relative yields of spin-dependent reaction pathways [6]. Because the downstream chemistry is influenced by the spin state, even weak magnetic fields may modulate protein activity and, in turn, biological function.

This phenomenon has been studied most extensively in cryptochromes, as well as in a small number of other flavoproteins, including photolyases and LOV-domain-containing flavoproteins [6–8]. However, the wide variety of biological processes affected by weak magnetic fields in a range of cells and organisms motivates a hypothesis that this effect may be found more widely among flavoproteins, and possibly within other classes of proteins as well as other biomolecules. This motivates the development of techniques to rapidly and reliably screen flavoproteins for magnetosensitivity.

One reliable readout for magnetosensitivity in flavoproteins is their spin-dependent fluorescence emission, in which fluorescence intensity changes in response to changing magnetic field conditions (also known as a magnetic field effect, or MFE). Singlet states can recombine more readily than triplet states back to the ground state; electrons can then be re-excited and emit fluorescence. Therefore, on average, fluorescence intensity is thought to be a good proxy for singlet yields. Recent work has demonstrated that a LOV domain can be engineered through directed evolution both to enhance the MFE and to change the kinetics of the response [9, 10]. Fluorescence is also an attractive approach for screening flavoproteins for magnetosensitivity because it in principle does not require developing a separate activity assay for each flavoprotein.

However, fluorescence-based MFEs in natural magnetosensitive proteins are often small and can be difficult to detect reproducibly. For example, the most well-studied protein in this space, cryptochrome, has been demonstrated to have an MFE (measured as a percent change in fluorescence upon application of an external magnetic field) of only 0.05–0.7%, depending on the imaging conditions [11, 12]. In these studies, furthermore, illumination intensity is not always reported, and the impact of illumination intensity on MFE magnitude is mostly not considered. In this work, we demonstrate that sufficient illumination intensity is critical to detect an MFE in the fluorescence of flavoproteins. Through ordinary differential equations (ODE) kinetics modeling, we demonstrate that photoexcitation rates have a strong influence on fluorescence contrast between singlet and triplet states and on the kinetics of fluorescence changes upon changing magnetic field conditions. As we will demonstrate, observing spin-state-dependent fluorescence contrast requires illumination intensities high enough that photoexcitation is not the rate-limiting step.

## 2. Methods and Results

Here, we propose a four-state ODE model with the following states: a flavin in the ground electronic state (state *F*); the flavin in the excited electronic state (state *F*\*); a radical pair (RP) in the singlet state (RP_*S*_); and a radical pair in the triplet state (RP_*T*_). The rate of excitation of *F* to *F*\* is given by *k*_exc_, whereas the rate of relaxation of *F*\* down to *F*, our observable in experiments and the only fluorescent transition, is given by *k*_f_. The rate of electron transfer to form a radical pair is given by *k*_ET_. The electron donor that participates in electron transfer is assumed to be intramolecular (within the flavoprotein), so RP formation is assumed to be unimolecular. The full model is depicted in Fig. 1, with all the relevant model parameters defined in Table I.

**Fig. 1.**
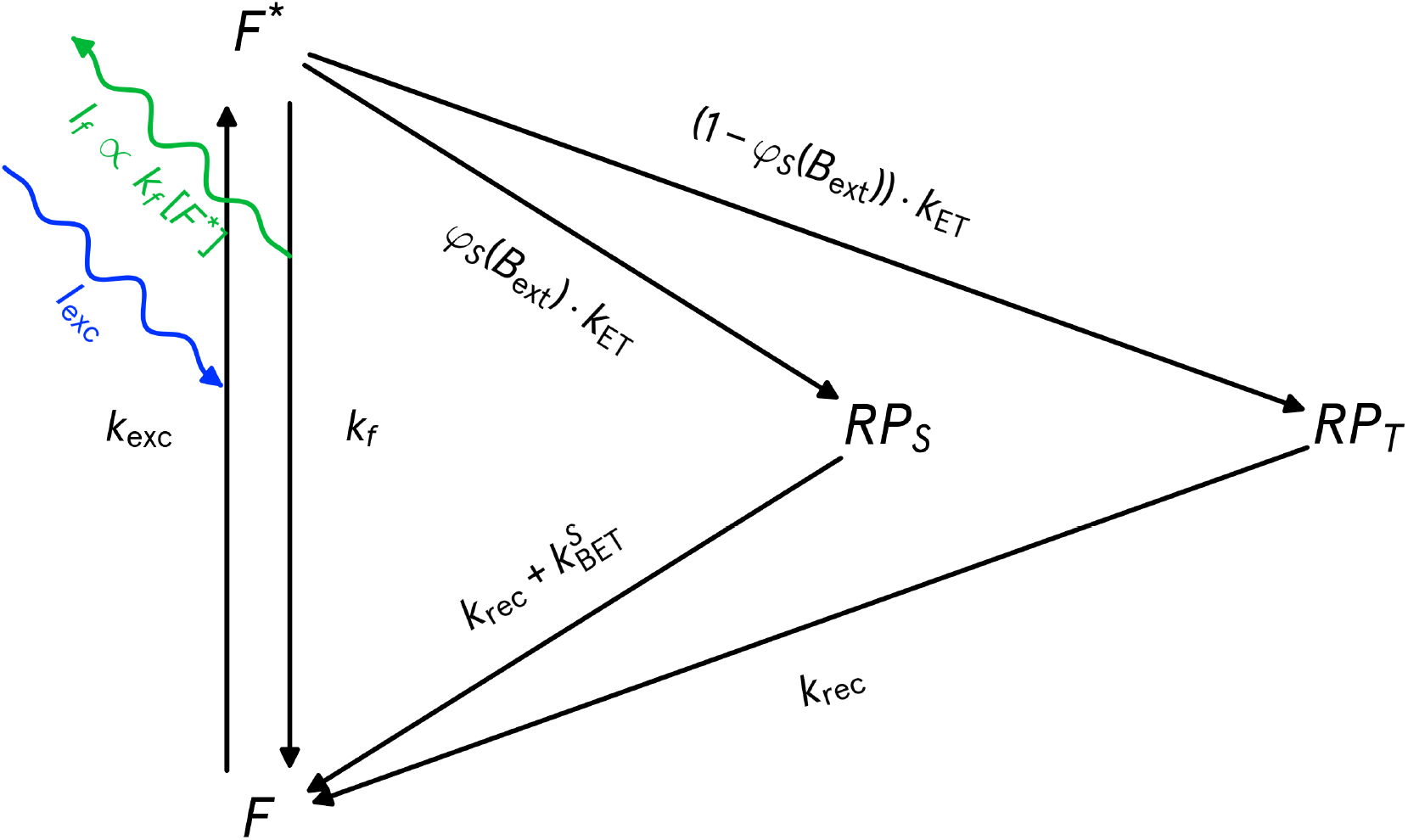
4-state ODE kinetic model of the radical pair mechanism, connecting spin chemistry to fluorescence readout. Schematic of the 4-state ODE model. See Table I for definitions.

**Table I.** States and parameters used in the kinetic model, and their origins.

| States | Meaning | Initial Value | Comment |
| --- | --- | --- | --- |
| $F$ | Flavin (ground state) | 1 | Assumes all flavin starts at the ground state |
| $F^*$ | Flavin (excited state) | 0 | |
| $RP_S$ | Radical pair (singlet) | 0 | |
| $RP_T$ | Radical pair (triplet) | 0 | |
| Conserved Quantity | Meaning | Total Value | Comment |
| $F_{\text{tot}}$ | Total flavin population | 1 | $F_{\text{tot}} = F + F^* + RP_S + RP_T$ |
| Independent Parameters | Meaning | Default Value | Comment |
| $k_{\text{exc}}$ | Excitation rate | $0.6 \text{ s}^{-1}$ | Corresponds to illumination intensity ($I_{\text{exc}}$) of $5 \text{ mW/cm}^2$, based on measurements of illumination from our own custom magnetofluorescence screening system. See $I_{\text{exc}}$ entry for equation [10] comparing $k_{\text{exc}}$ and $I_{\text{exc}}$ |
| $k_f$ | Fluorescence rate | $2 \times 10^8 \text{ s}^{-1}$ | Based off of fluorescent lifetime of flavin-binding fluorescent proteins between 3.17 and 5.7 ns [14] |
| $k_{\text{ET}}$ | Rate of electron transfer | $10^{10} \text{ s}^{-1}$ | From [11] |
| $k_{\text{BET}}^S$ | Singlet-specific back electron transfer | $2.0 \times 10^6 \text{ s}^{-1}$ | From [11] |
| $k_{\text{rec}}$ | Spin-independent ground state recovery via slow termination reactions | $0.01 \text{ s}^{-1}$ | Abrahams *et al.* (2026) [10] demonstrate slow FMNH• (neutral semiquinone radical state) formation over minutes for MagLOV2 |
| $\varphi_S(B_{\text{ext}})$ | Singlet yield | 0.208 with $B_{\text{ext}}$ OFF;<br>0.167 with $B_{\text{ext}}$ ON | Based on singlet yields of two electrons and one nuclear spin for triplet-born radical pairs with Larmor frequency $\omega = 0^+$ ($B_{\text{ext}}$ OFF) and $\omega \rightarrow \infty$ ($B_{\text{ext}}$ ON) [13]. However, we do not know the actual singlet yields of a flavoprotein or its magnetic field dependence, so the present choice is somewhat arbitrary. A full quantum mechanical model is needed to predict singlet yield's dependence on magnetic fields. Note that the magnetic field dependence is encoded by changing the value of this parameter |
| $\varepsilon$ | Extinction coefficient | 13,728 M<sup>-1</sup>cm<sup>-1</sup> | $\varepsilon(447 \text{ nm}) = 13800 \text{ M}^{-1}\text{cm}^{-1}$ was measured for AsLOV2 protein [15]. This value was rescaled to the extinction coefficient at 450 nm, based on the riboflavin absorption spectrum [16] |
| $\lambda$ | Excitation wavelength | 450 nm | Approximate peak of flavin absorbance spectrum |

| Derived Parameters | Meaning | Equation | Comment |
| --- | --- | --- | --- |
| $I_{\text{exc}}$ | Illumination intensity | $I_{\text{exc}} = \frac{k_{\text{exc}} N_A h c}{1000 \ln(10) \lambda \varepsilon(\lambda)}$ | Excitation rate from photon flux: $k_{\text{exc}} = \sigma \Phi_{\text{photon}}$ with $\sigma = 1000 \ln(10) \varepsilon(\lambda) / N_A$, standard photophysics formula [17] |
| $I_{\text{f}}$ | Fluorescence intensity | $I_{\text{f}} = k_{\text{f}} [F^*]$ | Fluorescence intensity is proportional to transitions from flavin excited state ($F^*$) back to ground state ($F$) |
| $C$ | Singlet-triplet contrast | $C = \frac{I_{\text{f}}(\varphi_S=1) - I_{\text{f}}(\varphi_S=0)}{I_{\text{f}}(\varphi_S=1)}$ | The difference between fluorescence intensity for pure singlet radical pairs and pure triplet radical pairs, normalized to the fluorescence intensity of pure singlet radical pairs |

The probability that an RP results in a singlet state (RP_*S*_) is given as *φ*_*S*_, and the probability that it is formed in a triplet state is given as 1 − *φ*_*S*_. A full quantum model of the Hamiltonian, modeling the two electrons and nearby nuclear spins, would be required to calculate the singlet and triplet yield-dependence on magnetic field intensity [13]. However, for this simplified, classical model, *φ*_*S*_ is considered an input parameter to the model and is treated as the average proportion of time the radical pair spends in the singlet state. It is this parameter that is treated as sensitive to external magnetic fields (*B*_ext_). The RP can recombine back to *F* in a spin-dependent manner. Singlets can undergo back electron transfer 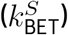, which recovers the ground state, while triplets cannot undergo this process because it is spin-prohibited. However, slower forward reactions involving protonation, deprotonation, and oxidation can also recover the *F* state in a spin-independent way, and these combined processes are described by the kinetic parameter *k*_rec_.

The following system of equations thus constitutes the model:

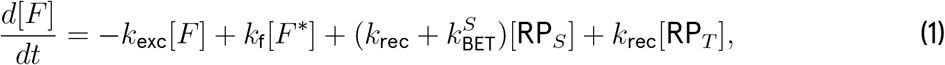

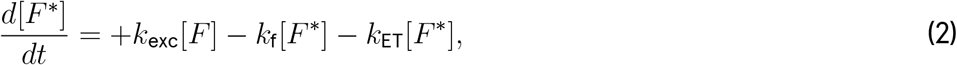

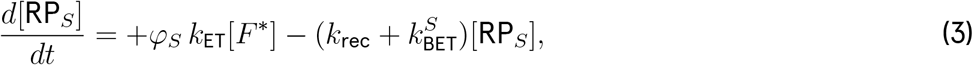

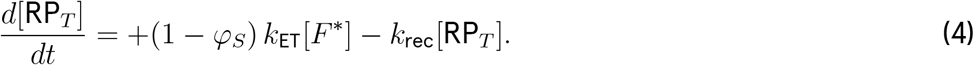

Furthermore, the total flavin concentration [*F*_tot_] is conserved:

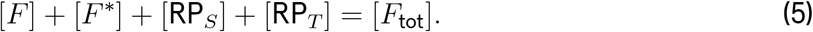

Fluorescence is the result of radiative relaxation from the excited *F*\* state back down to the ground state *F*, and fluorescence intensity is given by the equation:

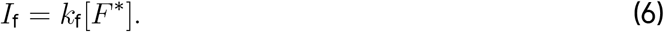

To relate the illumination intensity (*I*_exc_) to *k*_exc_, the following relationship holds:

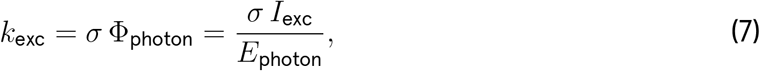

where *σ* is the single-molecule cross-section absorption area of the flavin molecule, Φ_photon_ is the photon flux, and *E*_photon_ is the energy of a photon. By substituting in the equation the energy of a photon, the following relationship is obtained:

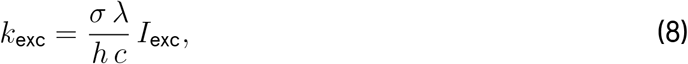

where *λ* is the wavelength of light, *h* is Planck’s constant, and *c* is the speed of light. The cross-section area of the molecule can be estimated using the relationship:

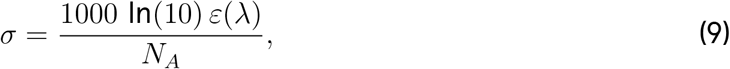

where *ε* is the extinction coefficient in M^*−*1^cm^*−*1^ and *N*_*A*_ is Avogadro’s number [17]. Plugging the formula for *σ* into the previous relationship and subsequently solving for *I*_exc_ results in an expression for *k*_exc_ in terms of *I*_exc_:

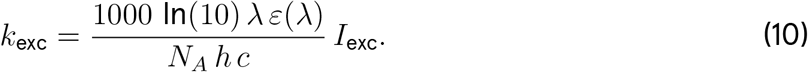

To screen proteins for effects of weak magnetic fields, which affect singlet yield, one could conveniently use fluorescence intensity as an experimental readout. To find sets of parameters for which fluorescence intensity can serve as a proxy measurement for singlet yield, we can calculate the fluorescence contrast, or the normalized difference in fluorescence intensity for simulations with pure singlet (*φ*_*S*_ = 1) and pure triplet (*φ*_*S*_ = 0) states:

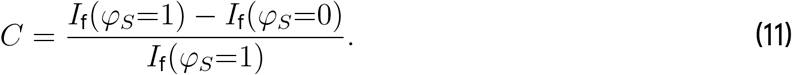

Contrast values close to 1 indicate parameter sets for which fluorescence can serve as a good proxy measurement for singlet activity, whereas values close to 0 indicate when changes in RP spin state would not result in a significant fluorescence intensity change. We computed the singlet-triplet fluorescence contrast over a range of excitation intensities and kinetic parameters (Fig. 2).

**Fig. 2.**
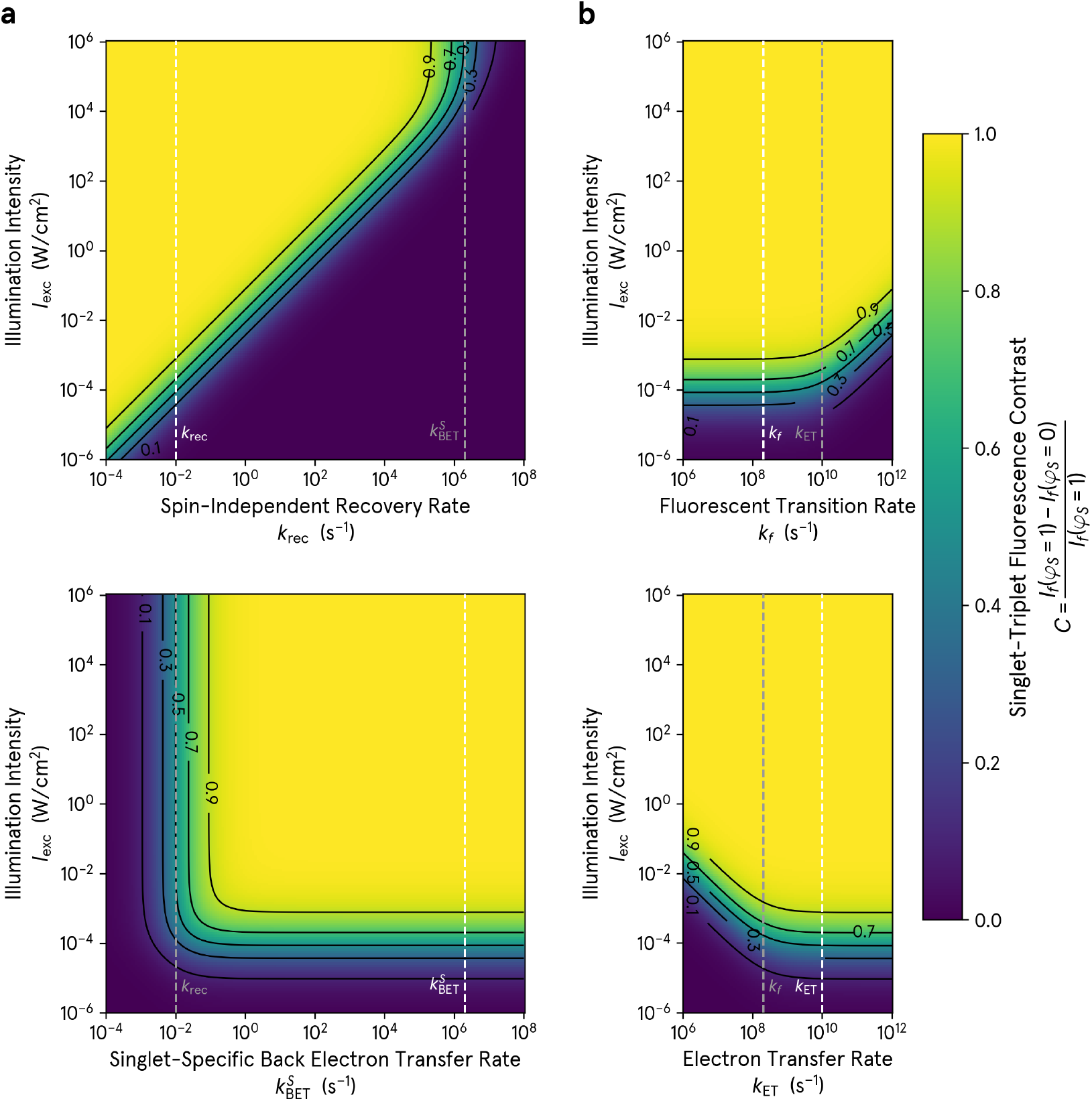
Parameter search for kinetic regimes in which fluorescence intensity is a good proxy for singlet yield indicates that the illumination intensity required for singlet-triplet fluorescence contrast is sensitive to the rate of spin-independent ground state recovery. Heat maps of fluorescence contrasts between pure singlet and pure triplet radical pairs for different combinations of illumination intensities (*I*_exc_) and the four kinetic parameters of the model: *k*_rec_ (**a**, top), 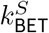 (**a**, bottom), *k*_f_ (**b**, top), and *k*_ET_ (**b**, bottom). Default parameters used in our simulation are shown with dashed lines. Around the default values, the illumination intensity required for fluorescence contrast of singlet and triplet states is most sensitive to *k*_rec_.

We find that, of the four kinetic rates, the illumination intensity *I*_exc_ needed to obtain high singlet-triplet fluorescence contrast was most sensitive to changes in *k*_rec_ (Fig. 2a). We find that the excitation intensity required to achieve high contrast scales approximately linearly with the spin-independent recombination rate *k*_rec_ (Fig. 2a). However, when 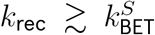,the contrast is strongly suppressed even at high excitation intensities, as spin-independent recombination dominates over spin-selective pathways, rendering the fluorescence effectively insensitive to the spin state (Fig. 2a). The photophysical rates impose a second condition (Fig. 2b): as long as *k*_f_ *< k*_ET_, most excited molecules proceed to a radical pair and the *I*_exc_ threshold for contrast is independent of *k*_f_. Only once *k*_f_ *> k*_ET_, where fluorescence begins to outcompete radical-pair formation, does the required *I*_exc_ depend on *k*_f_ or *k*_ET_. Together, these observations suggest that fluorescence contrast between singlet and triplet states is most sensitive to the slowest kinetic parameter in our model, namely, the spin-independent chemical reactions that recover the ground state fluorophore. **Thus, fluorescence is only a reliable proxy measurement for singlet yield when photoexcitation outcompetes spin-independent ground-state recovery processes**.

We then modeled the fluorescence response *I*_f_ over time to understand the relationship between the parameters of the model and the magnitude and kinetics of MFEs in the fluorescence output in response to changing magnetic field conditions (Fig. 3). A full quantum-mechanical treatment of the electron and nuclear spin dynamics and the hyperfine-driven singlet-triplet mixing during the superposition coherence time of the RP would be required to determine the exact dependence of *φ*_*S*_ on magnetic-field strength. In this classical model, however, *φ*_*S*_(*B*_ext_, OFF) and *φ*_*S*_(*B*_ext_, ON) were treated as input parameters, which switched instantaneously upon application or removal of *B*_ext_. In the plots of Fig. 3, the system was allowed to reach a steady state with *φ*_*S*_(*B*_ext_, OFF), and subsequently the value of *φ*_*S*_ was switched to *φ*_*S*_(*B*_ext_, ON) at *t* = 0, reflecting the application of an external magnetic field. For the chosen default parameters, fluorescence decreases after *t* = 0 (Fig. 3a); the sign of the MFE (decreasing as opposed to increasing) is due to the fact, explained below, that our chosen *φ*_*S*_(*B*_ext_, OFF) *> φ*_*S*_(*B*_ext_, ON). Running these simulations for different illumination intensities resulted in changes in both the magnitude of the response and the kinetics (Fig. 3b). The magnitude of the response increased with illumination before saturating, whereas its speed continued to rise as illumination increased.

**Fig. 3.**
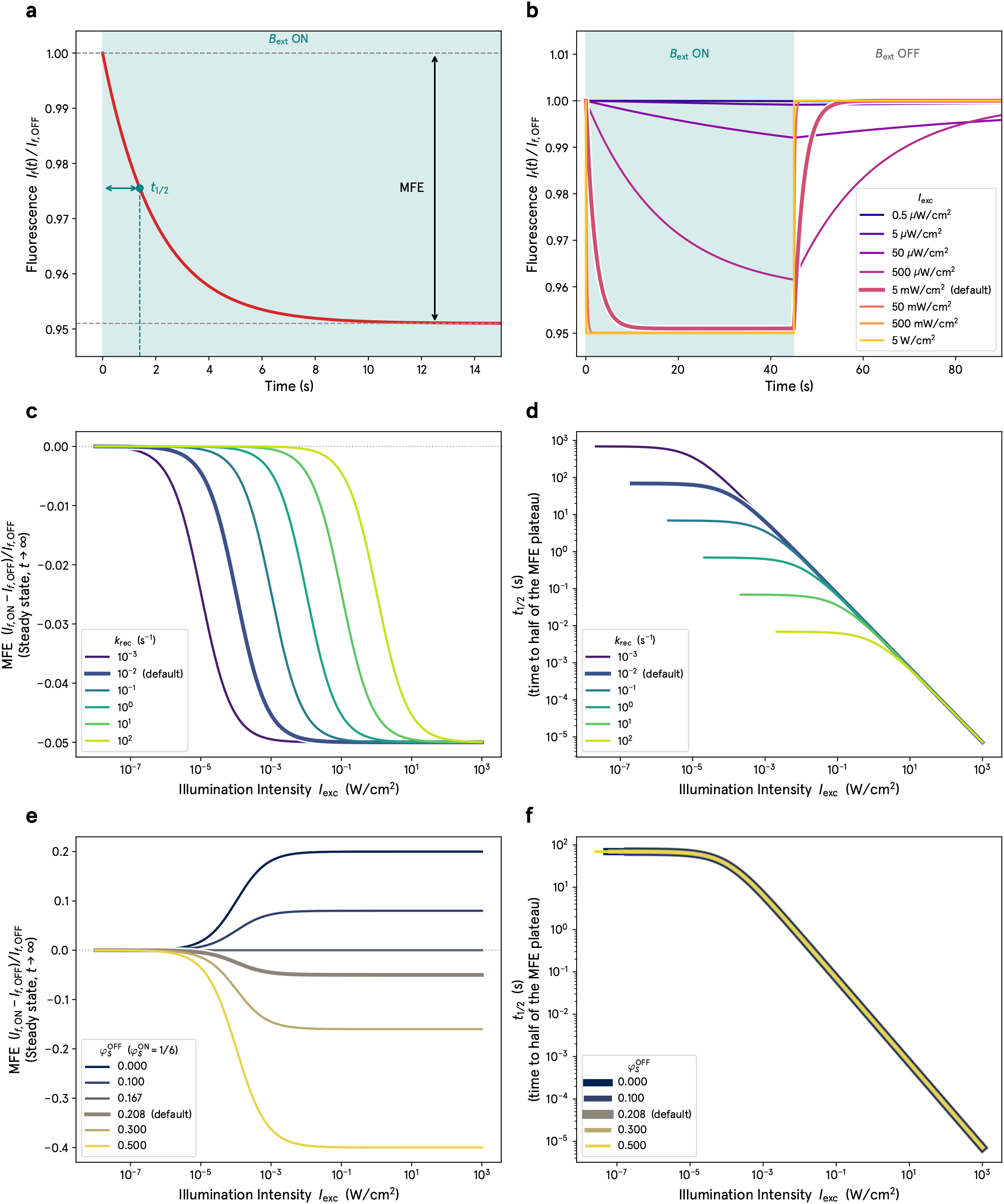
Illumination intensity affects both the magnitude and the kinetics of the fluorescence magnetic field effect, with spin-independent recovery and the field-induced change in singlet yield acting on distinct features of the response. **a)** Normalized modeled fluorescence traces 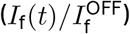 for the 4-state model with default parameters. An external magnetic field (*B*_ext_) is applied at *t* = 0, reflecting an instantaneous change in the value of the singlet yield (*φ*_*S*_). The magnetic field effect (MFE) is measured as the percent change in fluorescence upon application of *B*_ext_ after it has reached a steady state (as *t* → ∞), and *τ*_1/2_ is the observed time constant of the response. **b)** Normalized modeled fluorescence traces sweeping different values of the illumination intensity (*I*_exc_). The value of *φ*_*S*_ switches instantaneously from that for magnetic field OFF to magnetic field ON at *t* = 0 s and back to OFF at *t* = 45 s. The magnitude and kinetics of the response are both enhanced as *I*_exc_ increases. **c)** MFE vs *I*_exc_ plots, sweeping different values of *k*_rec_. The MFE vanishes at low *I*_exc_ and saturates at high *I*_exc_. Increasing *k*_rec_ shifts the inflection point to higher values of *I*_exc_, meaning that photocycles with faster spin-independent ground state recovery require stronger illumination for an MFE to be experimentally observable. **d)** *τ*_1/2_ vs. *I*_exc_ plots sweeping different values of *k*_rec_. The value of *τ*_1/2_ decreases as *I*_exc_ increases, and the kinetics at lower *I*_exc_ depend on *k*_rec_, while at higher *I*_exc_ it loses its dependence on *k*_rec_. **e)** MFE vs. *I*_exc_ curves, sweeping *φ*_*S*_(*B*_ext_, OFF), with *φ*_*S*_(*B*_ext_, ON) fixed at 0.167. Varying *φ*_*S*_(*B*_ext_, OFF) sets the sign and magnitude of the MFE plateau (positive when *φ*_*S*_(*B*_ext_, OFF) *< φ*_*S*_(*B*_ext_, ON), negative when greater). **f)** *τ*_1/2_ vs. *I*_exc_ curves, sweeping different values of *φ*_*S*_(*B*_ext_, OFF), with *φ*_*S*_(*B*_ext_, ON) fixed at 0.167. The kinetics of the response are independent of the field-induced singlet yield change.

From time traces of *I*_f_, we extracted two parameters: the magnitude of the change in fluorescence after having reached steady state (i.e., the magnitude of the MFE) and the observed time constant (*τ*_1/2_), the time for the fluorescence to change by half of the total MFE (Fig. 3a). We then plotted MFE and *τ*_1/2_ vs *I*_exc_ for varying values of *k*_rec_ and *φ*_*S*_(*B*_ext_, OFF), both by analytically solving the ODEs (Fig. 3, see Supplemental Results and Methods for analytical solutions) and by solving them numerically (SI Figure 1). MFE vs *I*_exc_ plots demonstrate that MFEs vanish at very low illumination intensities and plateau at higher illumination intensities (Fig. 3c). Increasing *k*_rec_ shifts the inflection point to higher illumination intensities, consistent with the observation that illumination intensities needed to get singlet-triplet fluorescence contrast scales with *k*_rec_ (Fig. 2). However, the amplitude of the MFE plateau at high values of *I*_exc_ was not affected by *k*_rec_. The time constant *τ*_1/2_, on the other hand, is determined analytically by ln(2)/*k*_rec_ at low illumination intensities. In contrast, at higher illumination intensities, *τ*_1/2_ starts to decrease with increasing *I*_exc_ and stops being dependent on the value of *k*_rec_, reflecting enhanced kinetics at high illumination intensities (Fig. 3d). Varying *φ*_*S*_(*B*_ext_, OFF) does not shift the inflection point of the MFE as a function of illumination intensity, but does affect the MFE’s final value plateau, with positive MFEs when *φ*_*S*_(*B*_ext_, OFF) *< φ*_*S*_(*B*_ext_, ON) and negative MFEs when *φ*_*S*_(*B*_ext_, OFF) *> φ*_*S*_(*B*_ext_, ON) (Fig. 3e). Varying *φ*_*S*_(*B*_ext_, OFF) has no effect on the kinetics of the response, as *τ*_1/2_ is independent of singlet yield changes upon application of an external magnetic field (Fig. 3f).

## 3. Discussion

This kinetic model demonstrates that having sufficient illumination intensity is critical for observing MFEs through a fluorescence readout. Fluorescence intensity integrated over the spin coherence times is often thought to be an unconditional proxy for singlet yield. Singlet radical pairs recombine through back electron transfer to the ground state, which can then be excited and fluoresce, while triplet states cannot and instead form forward photoproducts, which only reform the ground state through relatively slow protonation-deprotonation and redox reactions [10]. Our model demonstrates, however, that fluorescence is only a good proxy for singlet yield if the recombination step from the radical pair back down to ground state, namely, the only spin-dependent step, is the rate-limiting step. On the other hand, if illumination intensity is too low and excitation becomes the rate-limiting step, there will be a reduced fluorescence contrast.

Recent work with the engineering of the magnetosensitive fluorescent protein MagLOV demonstrated that both the magnitude and the kinetics of the magnetic field-sensitive fluorescence change can be selected for via directed evolution [9, 10]. Our kinetic model demonstrates that the magnitude of the MFE, for a given illumination intensity, can be altered either by altering the magnetodependence of the singlet yield (for example, by altering the hyperfine interactions conferred by the nuclear spins in proximity to the radicals), or by slowing down the spin-independent pathways that recover ground state flavin (for example through protonation-deprotonation reactions and slow redox processes) (Fig. 3).

For example, in the study by Abrahams *et al*., it was demonstrated that MagLOV2 forms the neutral semiquinone state (FMNH•, with the center dot symbolizing a radical form) very slowly, on the order of minutes [10]. FMNH•, acting as an intermediate between the RP and *F* in our model, must then oxidize and deprotonate to recover the quinone ground state (FMN, or *F* in our model) [10]. Because screening for magnetosensitive fluorescence responses during the development of MagLOV was likely done at a relatively low intensity, it is possible that evolved protein variants that slowed down the photocycle and thus allowed for a greater MFE in their fluorescence emission at low illumination intensities were screened for. Likewise, the authors demonstrated that a single mutation from MagLOV2 resulted in MagLOVf, which has faster kinetics but a smaller MFE. This is consistent with our model, which suggests that without sufficient illumination intensity, there is a tradeoff between the kinetics of the process and the magnitude of the MFE.

Previous work by Kattnig *et al*. demonstrated that MFEs on flavin photoreactions can be amplified chemically by slow radical termination reactions, and that this amplification was stronger at lower illumination intensities [11]. They showed that the slow radical-termination reactions downstream of the radical pair can substantially amplify or attenuate the observed MFE, depending on the ratio of the donor- and acceptor-radical termination rates. Our results make a complementary point at the level of intensity dependence: the same class of slow spin-independent recovery reactions, collapsed here into a single effective rate *k*_rec_, sets the illumination intensity required to observe an MFE via fluorescence intensity contrast at all. Together, these findings identify the slow termination and recovery chemistry as a primary control knob for magnetosensitivity alongside the spin-active steps, with direct implications for engineering MFE-based magnetosensitive fluorescent proteins: tuning the chemistry that returns radicals to the ground state after the radical pair loses coherence is as consequential as tuning the chemistry that forms the radical pair.

These results suggest that flavoproteins that do not exhibit an MFE in their fluorescence may nevertheless be magnetosensitive because the experimental results are setup-dependent. In these cases, the spin-independent recovery of the flavin cofactor may be too fast to get appreciable spin-dependent fluorescence contrast at reasonable illumination intensities, yet there may still be magnetic field-dependence of the singlet yield and thus downstream chemistry. **Therefore, any system designed to screen flavoproteins for magnetosensitivity via fluorescence needs to have sufficient illumination intensity so that spin-dependent steps are rate-limiting**. The lack of fluorescence contrast at low illumination intensities is due to the fundamental rates of the photocycle and is not a matter of the detector sensitivity or signal-to-noise ratio. When screening for magnetosensitivity for a particular set of proteins, it may be necessary to utilize many different illumination intensities, to balance increased singlet-triplet fluorescence contrast with photobleaching.

## 4. Conclusion and Next Steps

We have presented here a four-state ODE model to study how illumination intensity relates to fluorescence contrast between singlet and triplet pathways in chemical reactions governed by the radical pair mechanism within flavoproteins. The model demonstrates that the illumination intensity required to use fluorescence intensity meaningfully as a proxy for singlet yield depends on the rate of spin-independent processes that recover the oxidized ground state. Faster processes demand a higher illumination, which is critical for observing MFEs in flavoproteins via a fluorescent readout. This observation implies that screening setups that use fluorescence intensity as a readout may miss magnetosensitive processes if the completion of the photocycle is too fast relative to the rate of excitation from illumination. It also suggests that screening setups for magnetosensitivity with low illumination intensity may actually be screening for slower photocycles, not only large changes in singlet yield upon exposure to an external magnetic field. **It is thus critical that researchers in this scientific field start reporting illumination intensities when demonstrating MFEs**. In addition, better understanding the spin-independent processes that allow for ground state flavin recovery and how to modulate their kinetics would allow us to better rationally design proteins that show magnetosensitive fluorescence at low illumination intensities. Furthermore, the kinetics of MFEs are closely related to illumination intensity, and there may be tradeoffs between speed and amplitude for applications in which there are limitations on illumination intensity. Future steps would be to incorporate a full quantum-mechanical description of the spin-physics into the model. The findings here also motivate the development of a high-intensity magnetofluorescence screening system, so that chances to operate in the regime in which fluorescence intensity is a good proxy for singlet yield are increased.

## 5. Author Contributions

Following the CRediT taxonomy [18]:

1. Brian L. Ross: conceptualization; data curation; formal analysis; investigation; methodology; software; validation; visualization; and writing (original draft)
2. Alessandro Lodesani: conceptualization; funding acquisition; methodology; and writing (review and editing)
3. Clarice D. Aiello: conceptualization; funding acquisition; validation; and writing (review and editing)

## 6. Acknowledgements

We thank Dr. Jonathan R. Woodward for scientific discussions that were crucial in the conception of this work. We would like to thank Morgan Sosa for her assistance with formatting, revising, and editing the manuscript, and we thank Michael Montague and Venkatesh Sridharan for critically reading the manuscript and providing helpful feedback.

The [Quantum Biology Institute](http://quantumbiology.org/) is a California non-profit 501(c)(3) focused research organization that performs basic research underpinning the quantum biology field in an open-science fashion.

## Supplementary Information

**Figure S1.**
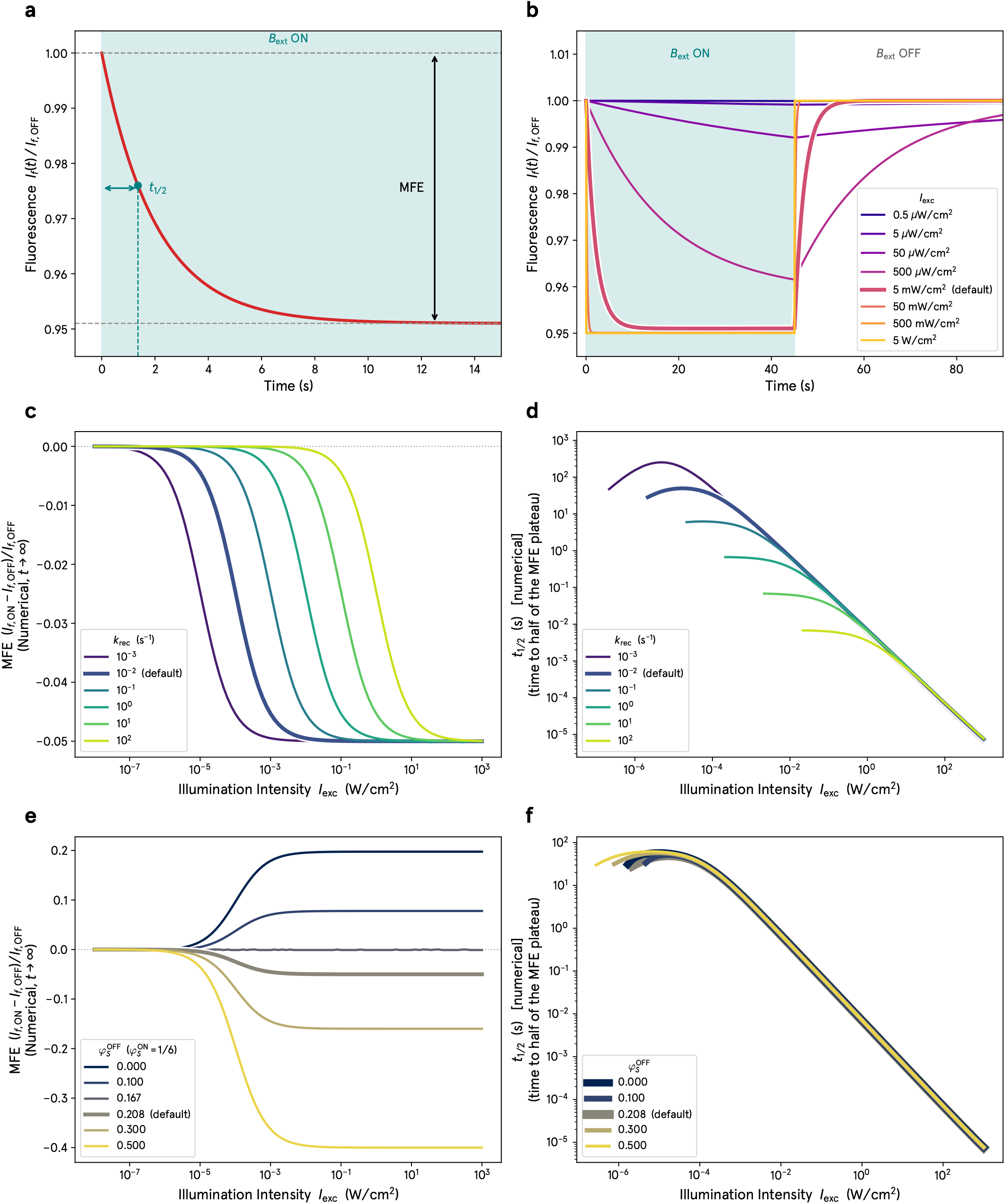
Solving the ODE model with numerical methods is consistent with the results from the explicit solution (shown in the main text Figure 3). Shown here is the same data as Figure 3, but solved using numerical methods. In these simulations, the model is run with default values (unless otherwise stated) first with the singlet yield value for the external magnetic field OFF until it reaches steady state (*φ*_*S*_(*B*_ext_, OFF)). Upon reaching steady state, the singlet yield value is instantaneously changed to that of the external magnetic field ON (*φ*_*S*_(*B*_ext_, ON)). **a)** Normalized modeled fluorescence traces 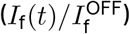 for the 4-state model with default parameters. An external magnetic field (*B*_ext_) is applied at *t* = 0, reflecting an instantaneous change in the value of the singlet yield (*φ*_*S*_). The magnetic field effect (MFE) is measured as the percent change in fluorescence upon application of *B*_ext_ after it has reached a steady state (as *t* → ∞), and *τ*_1/2_ is the observed time constant of the response. **b)** Normalized modeled fluorescence traces sweeping different values of the illumination intensity (*I*_exc_). The value of *φ*_*S*_ switches instantaneously from that for magnetic field OFF to magnetic field ON at *t* = 0 s and back to OFF at *t* = 45 s. The magnitude and kinetics of the response are both enhanced as *I*_exc_ increases. **c)** MFE vs. *I*_exc_ plots, sweeping different values of *k*_rec_. The MFE vanishes at low *I*_exc_ and saturates at high *I*_exc_. Increasing *k*_rec_ shifts the inflection point to higher values of *I*_exc_, meaning that photocycles with faster spin-independent ground state recovery require stronger illumination for an MFE to be experimentally observable. **d)** *τ*_1/2_ vs. *I*_exc_ plots sweeping different values of *k*_rec_. The value of *τ*_1/2_ decreases as *I*_exc_ increases, and the kinetics at lower *I*_exc_ depend on *k*_rec_, while at higher *I*_exc_ it loses its dependence on *k*_rec_. **e)** MFE vs. *I*_exc_ curves, sweeping *φ*_*S*_(*B*_ext_, OFF), with *φ*_*S*_(*B*_ext_, ON) fixed at 0.167. Varying *φ*_*S*_(*B*_ext_, OFF) sets the sign and magnitude of the MFE plateau (positive when *φ*_*S*_(*B*_ext_, OFF) *< φ*_*S*_(*B*_ext_, ON), negative when greater). **f)** *τ*_1/2_ vs *I*_exc_ curves, sweeping different values of *φ*_*S*_(*B*_ext_, OFF), with *φ*_*S*_(*B*_ext_, ON) fixed at 0.167. The kinetics of the response are independent of the field-induced singlet yield change.

## SI1. Methods and Results: Explicit Solution to the ODE Kinetic Model

The ODE model has four states: the ground state *F*, the excited state *F*\*, and the singlet and triplet radical pairs, RP_*S*_ and RP_*T*_, respectively. They obey the conservation law *F* + *F*\* + RP_*S*_ + RP_*T*_ = *F*_tot_, and they are described by the following set of ODEs:

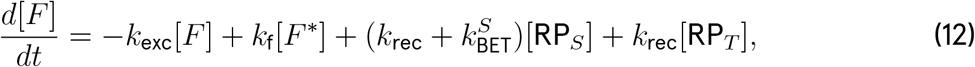

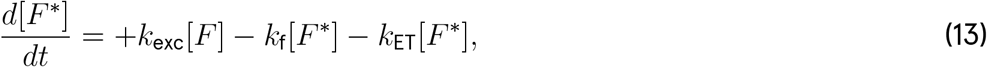

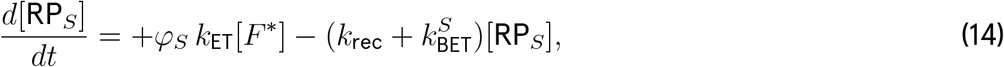

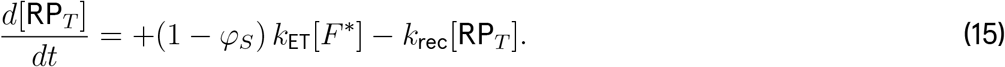

### Steady state explicit

To find the steady state explicit solution, we set the time derivatives to zero, which gives the following expressions:

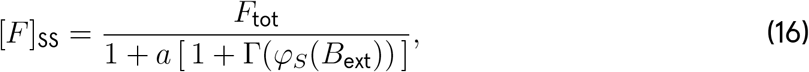

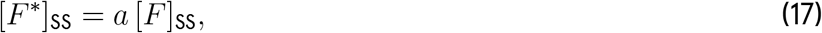

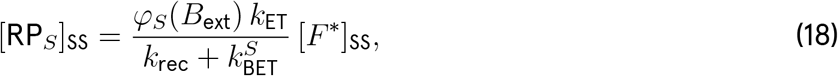

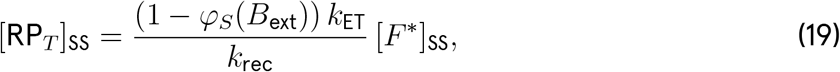

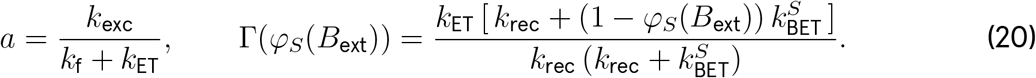

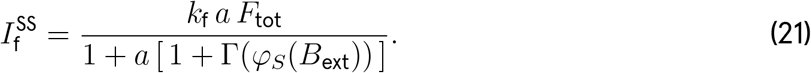

### Magnitude of the magnetic field effect

The magnitude of the magnetic field effect is given as the change in fluorescent intensity upon turning on the external magnetic field and reaching steady state:

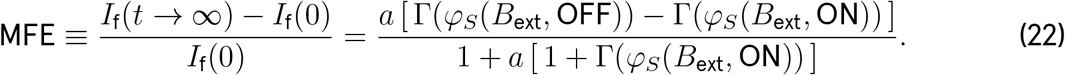

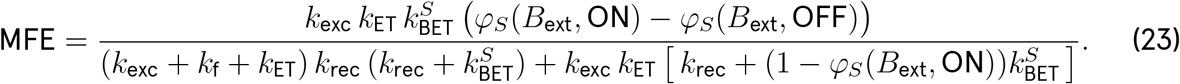

### Time-dependent explicit results and kinetics

Using the conservation law to remove *F* leaves a 3-state linear system, for which three relaxation rates *λ*_1_, *λ*_2_, *λ*_3_ are solutions to the characteristic equation:

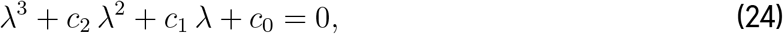

Where

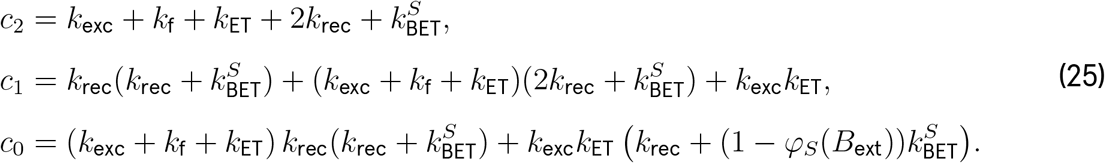

This can be solved explicitly in closed form using Cardano’s formula, but the resulting expressions are unwieldy. However, because the three roots are widely separated by several orders of magnitude, we can approximate the three eigenvalues, two fast relaxation modes and one slow mode, using either Vieta’s formulas or the method of dominant balance. The roots are thus given approximately by the following:

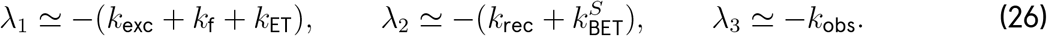

The first two rates, representing excited-state turnover and singlet-pair recombination respectively, are relatively fast. Thus the observed kinetics is dominated by the third slowest rate given by the following equation:

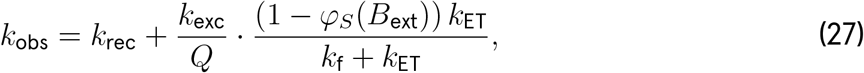

Where

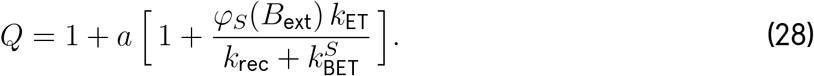

The half-time *τ*_1/2_ can be approximated as

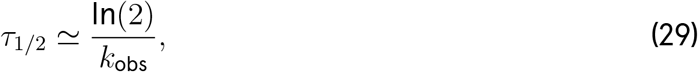

which can be written as a single expression:

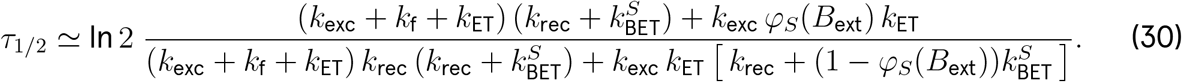

When excitation is slow relative to the depopulation of the excited state (*k*_exc_ ≪ *k*_ET_ + *k*_f_), then this expression simplifies to the following:

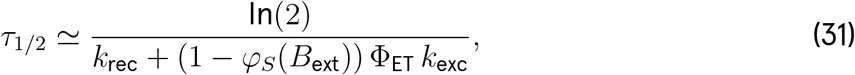

where Φ_ET_ is the electron transfer quantum yield,

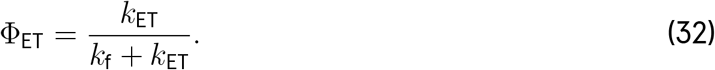

## References

[1] F. Bertagna, R. Lewis, S. R. P. Silva, J. McFadden, and K. Jeevaratnam, Effects of electromagnetic fields on neuronal ion channels: a systematic review, Ann. N. Y. Acad. Sci. 1499, 82 (2021).

[2] V. N. Binhi and F. S. Prato, Biological effects of the hypomagnetic field: An analytical review of experiments and theories, PLoS One 12, e0179340 (2017).

[3] H. Wang and X. Zhang, Magnetic fields and reactive oxygen species, Int. J. Mol. Sci. 18, 2175 (2017).

[4] H. Zadeh-Haghighi and C. Simon, Magnetic field effects in biology from the perspective of the radical pair mechanism, J. R. Soc. Interface 19, 20220325 (2022).

[5] A. Lodesani, G. Anders, L. Bougas, T. Lins, D. Budker, P. Fierlinger, and C. D. Aiello, eak magnetic field effects in biology are measurable—accelerated Xenopus embryogenesis in the absence of the geomagnetic field, bioRxiv (2024).

[6] P. J. Hore and H. Mouritsen, The radical-pair mechanism of magnetoreception, Annu. Rev. Biophys. 45, 299 (2016).

[7] R. Wiltschko, C. Nießner, and W. Wiltschko, The magnetic compass of birds: The role of cryptochrome, Front. Physiol. 12, 667000 (2021).

[8] K. B. Henbest, K. Maeda, P. J. Hore, M. Joshi, A. Bacher, R. Bittl, S. Weber, C. R. Timmel, and E. Schleicher, Magnetic-field effect on the photoactivation reaction of Escherichia coli DNA photolyase, Proc. Natl. Acad. Sci. U.S.A. 105, 14395 (2008).

[9] R. F. Hayward, A. Rai, J. R. Lazzari-Dean, A. E. Y. T. Lefebvre, A. G. York, and M. Ingaramo, Magnetic control of the brightness of fluorescent proteins, Protein Magnetofluorescence project site, Andrew G. York Lab, Calico Life Sciences (preprint) (2023), accessed 22 Jan 2026.

[10] G. Abrahams, A. Štuhec, V. Spreng, R. Henry, I. Kempf, J. James, K. Sechkar, S. Stacey, V. Trelles-Fernandez, L. M. Antill, C. R. Timmel, J. J. Miller, M. Ingaramo, A. G. York, J.-P. Tetienne, and H. Steel, Quantum spin resonance in engineered proteins for multimodal sensing, Nature 649, 1172 (2026).

[11] D. R. Kattnig, E. W. Evans, V. Déjean, C. A. Dodson, M. I. Wallace, S. R. Mackenzie, C. R. Timmel, and P. J. Hore, Chemical amplification of magnetic field effects relevant to avian magnetoreception, Nat. Chem. 8, 384 (2016).

[12] V. Déjean, M. Konowalczyk, J. Gravell, M. J. Golesworthy, C. Gunn, N. Pompe, O. F. Vander Elst, K.-J. Tan, M. Oxborrow, D. G. A. L. Aarts, S. R. Mackenzie, and C. R. Timmel, Detection of magnetic field effects by confocal microscopy, Chem. Sci. 11, 7772 (2020).

[13] C. D. Aiello, B. L. Ross, A. Lodesani, and M. L. Sosa, A physicist-friendly primer on the Hamiltonian for quantum sensing in proteins: analytical expressions and insights for a toy model of the radical-pair mechanism, arXiv:2604.18608 (2026), arXiv:2604.18608.

[14] M. Wingen, J. Potzkei, S. Endres, G. Casini, C. Rupprecht, C. Fahlke, U. Krauss, K.-E. Jaeger, T. Drepper, and T. Gensch, The photophysics of LOV-based fluorescent proteins – new tools for cell biology, Photochem. Photobiol. Sci. 13, 875 (2014).

[15] M. Salomon, J. M. Christie, E. Knieb, U. Lempert, and W. R. Briggs, Photochemical and mutational analysis of the FMN-binding domains of the plant blue light receptor, phototropin, Biochemistry 39, 9401 (2000).

[16] Oregon Medical Laser Center, PhotochemCAD: [Riboflavin], https://omlc.org/spectra/PhotochemCAD/html/004.html (2026), accessed 29 May 2026.

[17] S. E. Braslavsky, Glossary of terms used in photochemistry, 3rd edition (IUPAC recommendations 2006), Pure and Applied Chemistry 79, 293 (2007).

[18] CRediT – Contributor Roles Taxonomy, https://credit.niso.org/, accessed: 2026.

